# Neuropsychiatric Disorder Subtyping Via Clustered Deep Learning Classifier Explanations

**DOI:** 10.1101/2022.12.14.520428

**Authors:** Charles A. Ellis, Robyn L. Miller, Vince D. Calhoun

**Affiliations:** Tri-institutional Center for Translational Research in Neuroimaging and Data Science: Georgia State University, Georgia Institute of Technology, Emory University, Atlanta, GA 30303 USA; Tri-institutional Center for Translational Research in Neuroimaging and Data Science: Georgia State University, Georgia Institute of Technology, Emory University, Atlanta, GA 30303 USA.

## Abstract

Identifying subtypes of neuropsychiatric disorders based on characteristics of their brain activity has tremendous potential to contribute to a better understanding of those disorders and to the development of new diagnostic and personalized treatment approaches. Many studies focused on neuropsychiatric disorders examine the interaction of brain networks over time using dynamic functional network connectivity (dFNC) extracted from resting-state functional magnetic resonance imaging data. Some of these studies involve the use of either deep learning classifiers or traditional clustering approaches, but usually not both. In this study, we present a novel approach for subtyping individuals with neuropsychiatric disorders within the context of schizophrenia (SZ). We train an explainable deep learning classifier to differentiate between dFNC data from individuals with SZ and controls, obtaining a test accuracy of 79%. We next make use of cross-validation to obtain robust average explanations for SZ training participants across folds, identifying 5 SZ subtypes that each differ from controls in a distinct manner and that have different degrees of symptom severity. These subtypes specifically differ from one another in their interaction between the visual network and the subcortical, sensorimotor, and auditory networks and between the cerebellar network and the cognitive control and subcortical networks. Additionally, there are statistically significant differences in negative symptom scores between the subtypes. It is our hope that the proposed novel subtyping approach will contribute to the improved understanding and characterization of SZ and other neuropsychiatric disorders.

## I. Introduction

The identification of novel neuropsychiatric disorder subtypes based on brain recordings has many potential benefits including enhancing the scientific understanding of disorders, enabling improved data-driven diagnosis that goes beyond symptoms [1], or enabling personalized treatment. In this study, we identify several subtypes of schizophrenia (SZ) using a novel approach that combines explainable deep learning classifiers and traditional clustering methods to analyze resting-state functional magnetic resonance imaging (rs-fMRI) dynamic functional network connectivity (FNC).

Relative to other modalities like EEG [1]–[3], MEG [4], or MRI [5], rs-fMRI has several advantages for neuropsychiatric disorder analysis. Although fMRI has lower temporal resolution relative to EEG and MEG, it has higher spatial resolution and localization, enabling a better understanding of brain network interaction. Relative to MRI, fMRI offers insight into brain dynamics that have been associated with a variety of neuropsychiatric disorders [6], [7]. While a handful of studies have analyzed whole brain rs-fMRI activity, the high dimensionality of the data paired with the frequent relatively low number of available study participants makes it easier to analyze extracted features. A common type of extracted feature is FNC. Two main varieties of FNC include static FNC (sFNC) and dynamic FNC (dFNC). Both types of FNC have been applied to study a variety of neurological and neuropsychiatric disorders [8]– [10] as well as cognitive functions [10], [11]. However, it has been shown that dFNC contains richer information than sFNC and can help identify effects of neuropsychiatric disorders that would be overlooked in an sFNC analysis [12].

Many SZ dFNC studies have involved deep learning or machine learning-based classification [13], [14]. These studies do not typically focus on gaining new insights into SZ. They often involve explainability for insight into the key spatial or temporal features differentiating individuals with SZ from HC. However, because they aggregate explanations on a class-level and because SZ is known to be highly heterogeneous, it is likely that they overlook important differences between individuals with SZ that could be highly relevant for diagnosis and for better understanding the disorder. Efforts involving SZ dFNC clustering have often been more focused on extracting novel insights [6], [7] or presented methods that could provide novel insights [15], [16]. However, dFNC clustering studies have generally not focused on SZ subtyping, and the small number of studies [17] that have focused on SZ subtyping have done so without really considering which brain networks most differentiate each subtype from SZ. It is possible that explainable deep learning approaches for SZ dFNC classification could be combined with clustering to identify novel SZ subtypes based on their differences with healthy individuals (HCs).

In this study, we present a novel approach, Subtyping via Cluster Explanations (SpiCE), that involves clustering the explainability results of a one-dimensional convolutional neural network (1D-CNN) trained to identify individuals with a neuropsychiatric disorder. We apply the approach within the context of SZ, identifying 5 SZ subtypes with different patterns of brain activity and significant differences in symptom severity. We characterize each subtype, identifying the brain networks that differentiate them from HCs.

## II. Methods

### A. Description of Data and Preprocessing

In this study, we used the Functional Imaging Biomedical Informatics Research Network (FBIRN) rs-fMRI dataset. It has been used in multiple clustering [17][16] and classification studies [6] and can be made available upon a request to the authors. The dataset has 160 healthy controls (HCs) and 151 individuals with SZ (SZs). Data was collected from the University of New Mexico, the University of Iowa, the University of North Carolina at Chapel Hill, Duke University, the University of Minnesota, the University of California at Los Angeles, the University of California at Irvine, and the University of California at San Francisco. Data collection protocols were approved by the institutional review boards of the various universities, and all participants gave written informed consent. One 3T General Electric and six 3T Siemens scanners were used for collection. An AC-PC aligned echo-planar imaging (EPI) sequence was used to collect T2*-weighted functional images with TR=2s, TE=30ms, flip angle=77°, voxel size=3.4×3.4×4mm^3^, slice gap=1mm, 162 frames, and 5:24 minutes.

Rigid body motion correction accounted for head motion, and preprocessing was performed with statistical parametric mapping (https://www.fil.ion.ucl.ac.uk/spm/). The data was spatially normalized to an EPI template in the MNI space. Data was resampled to 3×3×3 mm^3^ and smoothed with a Gaussian kernel with a 6 mm full width at half maximum. After preprocessing, we performed feature extraction. The Neuromark automatic independent component (IC) analysis pipeline with the Neuromark_fMRI_1.0 template of the GIFT toolbox was used to extract 53 ICs. The 53 ICs were assigned to 7 networks with 5 subcortical (SCN), 9 sensorimotor (SMN), 2 auditory (ADN), 9 visual (VSN), 7 default mode (DMN), 17 cognitive control (CCN), and 4 cerebellar (CBN) network nodes. FNC between two networks is represented as network/network (e.g., SCN/VSN). We used a tapered sliding window approach, calculating Pearson’s correlation between ICs at each step, to obtain dFNC. The tapered window was made from a rectangle with a 40s step size convolved with a Gaussian with a standard deviation of 3.

### B. Description of Model Development

We developed a 1D-CNN architecture. Prior to training, we feature-wise z-scored each participant separately and used 10-fold stratified shuffle split cross-validation with 80-10-10 ratios of training-validation-test data. We quadrupled the training set size via a data augmentation approach in which we generated three copies of the training data and added Gaussian noise with a mean of zero and varying standard deviation for each copy (0.7, 0.5, 0.2). We used a class-weighted categorical cross-entropy loss function to reduce the potential effects of any class imbalances. We also used an Adam optimizer with an initial learning rate of 0.001 that decreased by 50% after each 15 epochs that passed without a corresponding increase in validation accuracy. We trained the model for 75 epochs with a batch size of 50. Kaiming He normal initialization was used for all layers except the first convolutional layer, which used Glorot Uniform initialization. The model from the epoch with the highest validation accuracy in each fold was used in testing. Test accuracy (ACC), sensitivity (SENS), and specificity (SPEC) were evaluated across folds.

### C. Description of Explainability-based Subtyping Approach

We used the αβ-rule [18] of layer-wise relevance propagation (LRP) [19] for explainability. LRP typically distributes either positive or negative relevance to the features of an input sample that totals to around 1, where positive relevance indicates features that support a sample being classified as the class of interest and negative relevance indicates features that support a sample being classified as an alternative class. We used the αβ-rule (α = 1, β = 0) to filter out negative relevance and only output positive relevance.

After outputting the relevance for each SZ training sample, we scaled the relevance of each sample such that it equaled 100 and summed the relevance assigned to each dFNC feature for each sample. We then calculated the average per-feature relevance for each training sample across folds. For example, if participant x appeared in the training set for 9 of 10 folds, then the 9 sets of relevance values for participant x were averaged, forming a single average explanation for that participant. After averaging relevance for each participant across folds, we applied k-means clustering to assign the relevance distributions to clusters. We used a number of clusters ranging from 2 to 10 and used the elbow method to select the optimal number of clusters.

### D. Description of Subtype-Specific Analysis of Symptom Severity

After assigning participants to clusters (i.e., subtypes), we sought to identify any subtype-specific differences in symptom severity. As such, we performed Kruskal-Wallis H-tests comparing the Positive and Negative Syndrome Scale (PANSS) positive and negative scores for each subtype.

## III. Results and Discussion

In this section, we describe and discuss our model performance, explainable subtyping results, differences in symptom severity between subtypes, and analysis next steps and limitations.

### A. Model Performance

Our classifier had an ACC, SENS, and SPEC of 79.38 ± 5.45, 76.88 ± 10.84, and 81.88 ± 7.1, respectively. Performance was above chance-level across all metrics. SPEC was about 5% higher than SENS, and ACC was near 80%. Our architecture obtained higher performance than some existing dFNC SZ classifiers [17][14].

### B. Identification of SZ Subtypes

Figure 2 shows the relevance centroids for each identified subtype. Figure 3 shows the mean dFNC for each class, and Figure 4 shows the mean dFNC for each subtype. Subtypes 1 through 5 have 9, 58, 53, 11, and 20 participants, respectively. For the most part, the model tended to rely upon different domain pairs for differentiating each subtype. For subtype 1, the model most relied upon VSN/SCN, VSN/SMN, and CBN/SCN. Subtype 4 similarly relied upon VSN/SCN and VSN/SMN, but it did not rely upon CBN/SCN. Additionally, comparable to subtypes 1 and 4, subtype 5 also relied upon VSN/SMN and, to a lesser degree, CBN/SCN but not upon VSN/SCN. Subtype 5 classification also seemed to rely more upon CCN/CBN and VSN/ADN. Subtypes 2 and 3 contained a larger number of samples than the other subtypes. As such, their mean relevance tended to be slightly lower. Subtype 3 relevance tended to be highest in CBN/SCN. The key network pairs for subtype 2 tended to have higher mean relevance than those of subtype 3. In particular, CBN/SCN and SMN/CBN were most important for subtype 2. These findings indicate that SZ seems to mainly affect the interactions of the VSN with the SCN, SMN, and ADN and of the CBN with the CCN and SCN to varying degrees. Many of these interactions were also identified in previous SZ subtyping efforts with the FBIRN dataset [17].

**Figure 1.**
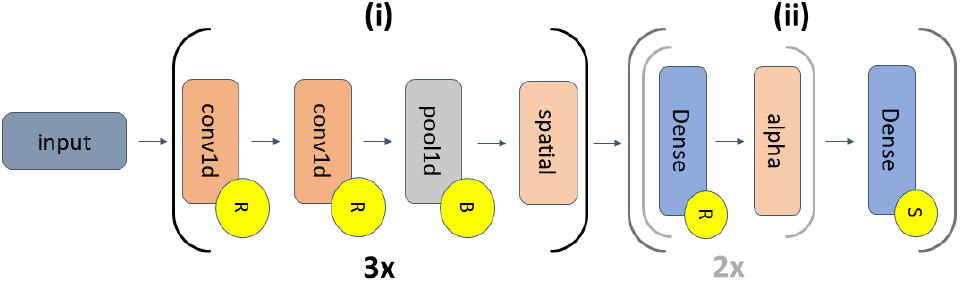
CNN Architecture. The model has feature extraction (i) and classification (ii) portions. (i) repeats three times. The grey inset within (ii) repeats twice. The first, second, and third convolutional (conv1d) layer (kernel size = 4) pairs have 16, 24, and 32 filters, respectively. Each pair is followed by a max pooling layer (pool size = 2) and spatial dropout (spatial, rates = 0.15, 0.25, and 0.3). Portion (ii) has 3 dense layers with 18, 14, and 2 nodes, respectively. The first two dense layers are followed by alpha dropout (alpha) with rates of 0.35 and 0.25. Yellow circles with “R”, “B”, and “S” show layers followed by ReLU activations, batch normalization, and softmax activations, respectively.

**Figure 2.**
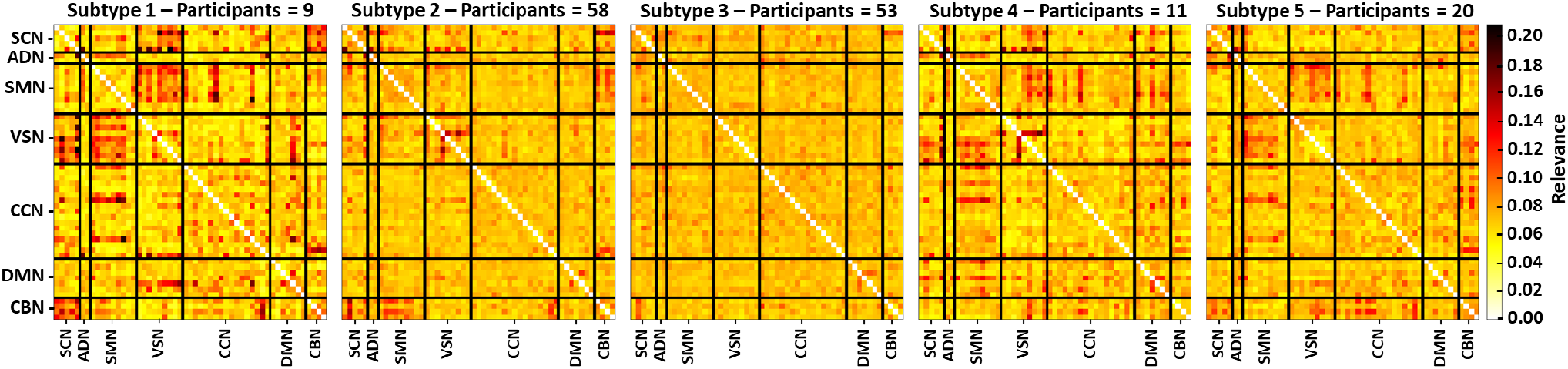
Centroid of Relevance for Each Subtype. The panel titles indicate their subtype and the number of samples assigned to that subtype. Each panel shares the same network labels on the y-axis, and the network labels for each panel are also on their x-axes. The panels share the same colormap and are arranged like a connectivity matrix to indicate the importance of the interaction of networks.

**Figure 3.**
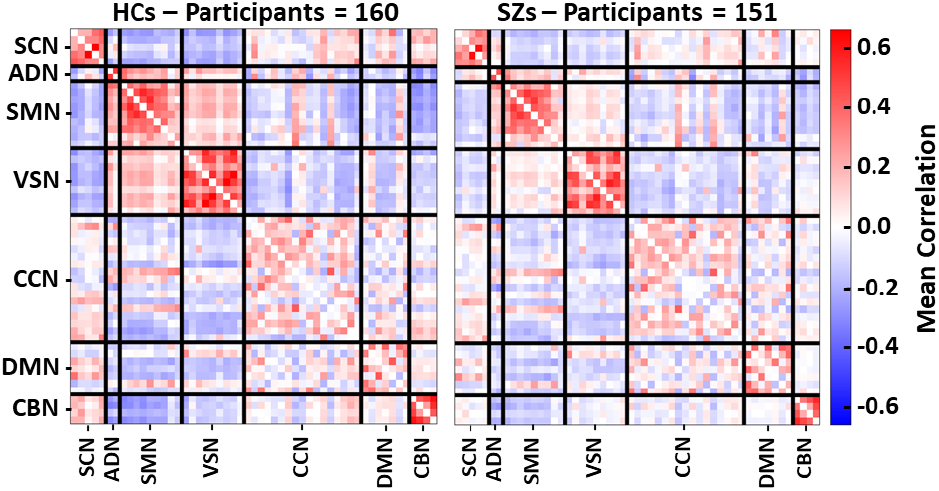
Mean dFNC for Each Class. The panel titles indicate their class and their number of samples. The panels share the same network labels on the y-axis, and the network labels for each panel are also on their x-axes. The panels share the same colormap.

**Figure 4.**
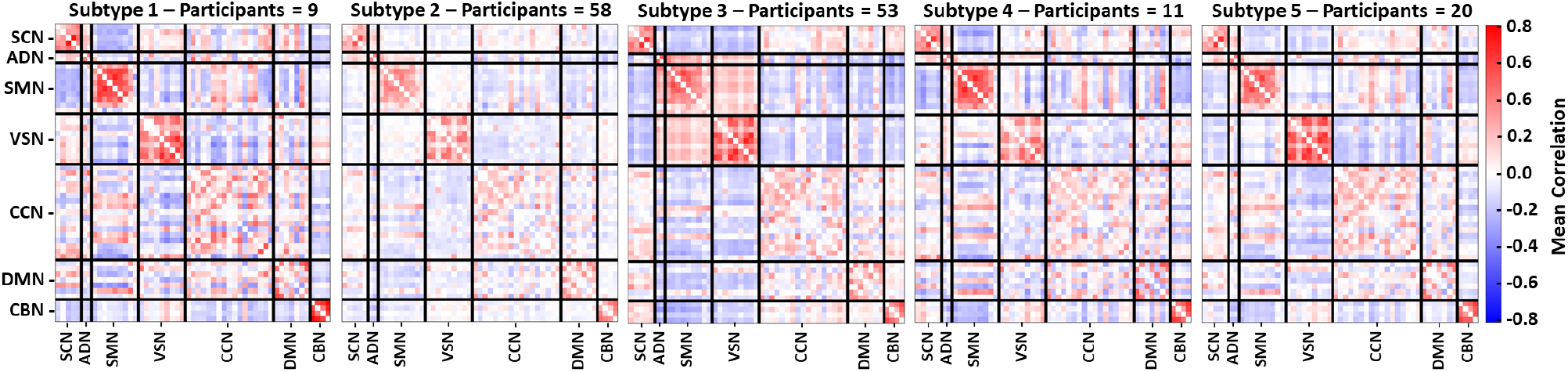
Mean dFNC for Each Subtype. The panel titles indicate their subtype and the number of samples assigned to that subtype. Each panel shares the same network labels on the y-axis, and the network labels for each panel are also on their x-axes. The panels share the same colormap.

Subtypes 1 and 4 tended to have more positive correlation for VSN/SCN relative to the other subtypes and HCs. In contrast, subtypes 1 and 4 tended to have more negative VSN/SMN correlation than other SZ subtypes, which further distinguished them from HCs [20]. Subtype 5 was unique in that VSN/ADN and CBN/CCN were particularly important for its classification and not for the other subtypes. However, those network pairs did not seem to have a systematic difference in mean connectivity relative to other subtypes, which could indicate that the effects of SZ upon those networks in SZ were more associated with how the dFNC varied over time. Subtypes 1, 2, and 5 had more negative CBN/SCN correlation than subtypes 3 and 4, while subtypes 3 and 4 seemed to have CBN/SCN activity more like HCs.

### C. Subtype-Specific Differences in Symptom Severity

Figure 5 shows the symptom severity across subtypes. The Kruskal-Wallis H-tests that we performed found a statistically significant difference in negative PANSS across subtypes but not in positive PANSS. Nevertheless, there are some visible differences in the distribution of positive PANSS across subtypes. Subtype 5, in general, had the highest symptom severity, which could result from the potential changes in VSN/ADN and CBN/CCN dynamics.

**Figure 5.**
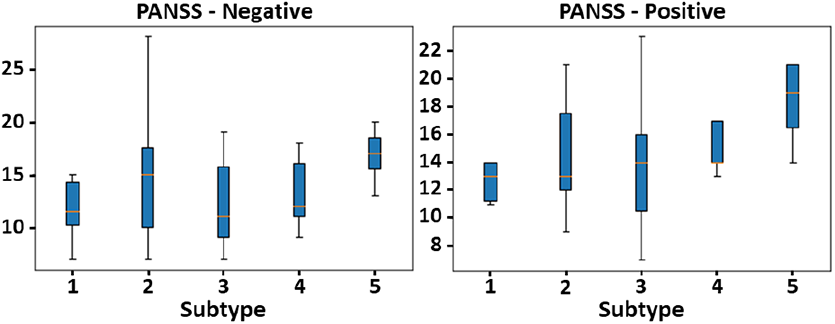
Symptom Severity for Each Subtype. The left and right panels show the negative and positive PANSS scores, respectively. The subtypes are arranged along the x-axis, and the y-axis for each panel show their respective symptom severity scores.

### D. Limitations and Next Steps

Our subtyping approach is unique to previous efforts that have sought to differentiate SZs from one another in that we sought to subtype SZs based upon the features that most differentiated them from HCs. While that is an important distinction, it also makes our subtyping highly dependent upon our classifier. Our results could be affected by (1) model performance or (2) even the random initialization of model weights during training. Our model test performance was quite high but not perfect. This is partially why we output explanations for the training set on which the model had near-perfect performance. We also sought to reduce the potential effects of random model initialization or other random effects of the training process by averaging the relevance for each participant across training sets from each fold. This should have provided a more robust estimate of the key network pairs for each participant. The approach we applied in this study could be applied to other neurological and psychiatric disorders to identify subtypes based on their differences from HCs. It could even be applied to identify subtypes based on the differences between one disorder and one or more other disorders with comparable symptoms.

## IV. Conclusion

Identifying neuropsychiatric disorder subtypes could contribute to a better understanding of those disorders and eventually to their improved diagnosis and treatment. In this study, we present a novel approach for subtyping neuropsychiatric disorders involving the clustering of explanations from a deep learning classifier trained on rs-fMRI dFNC data. We implement the approach within the context of SZ, identifying 5 subtypes with distinct patterns of brain network interaction and significant differences in symptom severity. We hope that our approach will contribute to the improved understanding of SZ and other neuropsychiatric disorders and to their eventual improved diagnosis and treatment.

## Acknowledgment

We thank those who collected the FBIRN dataset.

